# Competing interactions modulate the activity of Sgs1 during DNA end resection

**DOI:** 10.1101/515791

**Authors:** Kristina Kasaciunaite, Fergus Fettes, Maryna Levikova, Peter Daldrop, Petr Cejka, Ralf Seidel

**Affiliations:** Peter Debye Institute for Soft Matter Physics, Universität Leipzig, 04103 Leipzig, Germany; Institute of Molecular Cancer Research, University of Zurich, 8057 Zurich, Switzerland; Institute for Molecular Cell Biology, University of Münster, 48149 Münster, Germany; Institute for Research in Biomedicine, Università della Svizzera italiana, Faculty of Biomedical Sciences, CH-6500 Bellinzona, Switzerland; Department of Biology, Institute of Biochemistry, Eidgenössische Technische Hochschule (ETH) Zürich, Switzerland

**Keywords:** DNA repair, Dna2, homologous recombination, RecQ helicases, single-molecule

## Abstract

DNA double-strand break repair by homologous recombination employs long-range resection of the 5’ DNA ends at the break points. In *Saccharomyces cerevisiae*, this process can be performed by the RecQ helicase Sgs1 and the helicase-nuclease Dna2. Though functional interplay has been shown, it remains unclear whether and how the proteins cooperate on the molecular level. Here, we resolved the dynamics of DNA unwinding by Sgs1 at the single molecule level and investigated its regulation by Dna2, the single-stranded DNA binding protein RPA and the Top3-Rmi1 complex. We found that Dna2 modulates the velocity of Sgs1, indicating that during end resection the proteins form a physical complex and couple their activities. Sgs1 unwinds DNA and feeds single-stranded DNA to Dna2 for degradation. RPA is found to regulate the processivity and the affinity of Sgs1 to the DNA fork, while Top3-Rmi1 modulated the velocity of Sgs1. We think that the differential regulation of the Sgs1 activity by its protein partners is important to allow diverse cellular functions of Sgs1 during the maintenance of genome stability.

## Introduction

The genome of eukaryotic cells is constantly damaged by environmental factors, byproducts of the cellular metabolism as well as transactions of the DNA metabolism. Damages appear in a variety of forms, such as base lesions, cross-links between DNA strands or between DNA and proteins, as well as DNA single- and double-strand breaks (DSBs) [1]. To avoid genome instability [2], cells use a number of intricate mechanisms to repair DNA lesions. DSBs are usually repaired by either of two main mechanisms – non-homologous end joining (NHEJ) and homologous recombination (HR) [3–5].

HR uses in vegetative cells mostly genetic information stored in the sister chromatids in order to allow an error-free DSB repair [1]. This process is initiated by the resection of the 5’ DNA end at the break point, such that a 3’ overhang is created, which is immediately coated by the single strand DNA binding protein replication protein A (RPA). In *Saccharomyces cerevisiae* (budding yeast), long-range DNA end resection is driven either by the exonuclease Exo1 or by the helicase Sgs1 and the helicase/nuclease Dna2, which work in a synergistic manner [6]. In human cells, this conserved pathway is catalyzed by the Sgs1 homologs BLM or WRN together with human DNA2 [7–10]. Sgs1 is a processive 3’ to 5’ RecQ helicase [11,12]. In contrast, Dna2 possesses a highly processive and strictly unidirectional 5’ to 3’ motor activity [13,14], which likely functions as a ssDNA translocase in resection to facilitate the degradation of DNA unwound by Sgs1 [15,16]. DNA degradation during 5’ end resection is accomplished by the barrel-shaped nuclease domain of Dna2 that travels along DNA ahead of the helicase motor [17]. The action of Sgs1 and Dna2 motors with opposite polarity provides an intriguing similarity to the RecBCD complex that powers DNA end resection in *E.coli*. RecBCD uses the anti-parallel helicase activities of the RecD and RecB subunits for DNA unwinding and the nuclease activity of RecB for DNA degradation [18]. When reconstituting the DNA end resection reactions with the yeast proteins *in vitro*, a synergistic activity of Sgs1 and Dna2 was observed [19]. However, the underlying interplay of this interaction remains undefined. Both proteins fulfil a number of additional cellular functions, which suggests a much more flexible and dynamic situation compared to the stable RecBCD complex carrying out a single functionality. For example, Sgs1 is involved in other downstream processes of the HR pathway, including the dissolution of double Holliday junctions leading to non-crossovers as well as in the regulation of aberrant HR [20,21]. Dna2 is engaged in Okazaki fragment maturation [22], processing of stalled replication forks [10,23] as well as in checkpoint signaling [24].

Previous biochemical studies have revealed that Sgs1, RPA and Dna2 represent the minimal group of proteins that is able to reconstitute DNA end resection [19]. On the protein level, Sgs1 and RPA are known to interact via the large RPA70 subunit that binds to the N-terminal acidic region of Sgs1, which is located next to the helicase domain [25]. DNA unwinding by Sgs1 does not require RPA but was found to be stimulated by this protein [19]. In contrast, both nuclease and motor functions of Dna2 mandate cognate RPA [26,27]. In mice, both proteins interact through the N-terminal domain of the Rpa70 subunit and the alpha1 and OB folds of the Dna2 N-terminus [17]. RPA prevents 5’ end degradation by Dna2 and instead promotes 3’ end degradation, enforcing thus the correct polarity of DNA end resection [19,28].

Though functional synergies and physical protein-protein interactions have been identified, little is known, however, how Sgs1 and Dna2 as well as other protein partners cooperate at the molecular level, and which functional steps are affected by these interactions. In order to gain insight into these processes we studied DNA unwinding by Sgs1 using magnetic tweezers. This technique allowed us to monitor the DNA processing by Sgs1 and its physiological interaction partners - including RPA, Dna2 and the Top3-Rmi1 complex - in real-time on the single-molecule level. Our data reveal how the DNA unwinding activity of Sgs1 is differentially and dynamically modulated by its partners. Furthermore, we obtain evidence that Sgs1 and Dna2 form together with RPA a complex during the end resection reaction. Overall, our study helps to explain how the Sgs1 activities can be fine-tuned to achieve diverse cellular functions.

## Results

### DNA unwinding by Sgs1

To study DNA unwinding by Sgs1 and its interaction partners we employed a magnetic tweezers assay [13,14], in which a 6.6 kbp dsDNA molecule was bound at one end to a magnetic bead and on the other end to the surface of the fluidic cell of the magnetic tweezers setup (Fig 1A). A short 38 nucleotide (nt) gap with a 40 nt 5’ flap about 0.5 kbp away from the surface attachment supported the initiation of DNA unwinding by Sgs1 and Dna2. A pair of magnets above the fluidic cell was used to apply defined forces of 15 to 25 pN onto the magnetic bead and therefore to stretch the DNA. Video-microscopy was used to track the bead position and thus to monitor changes of the DNA length.

**Fig 1.**
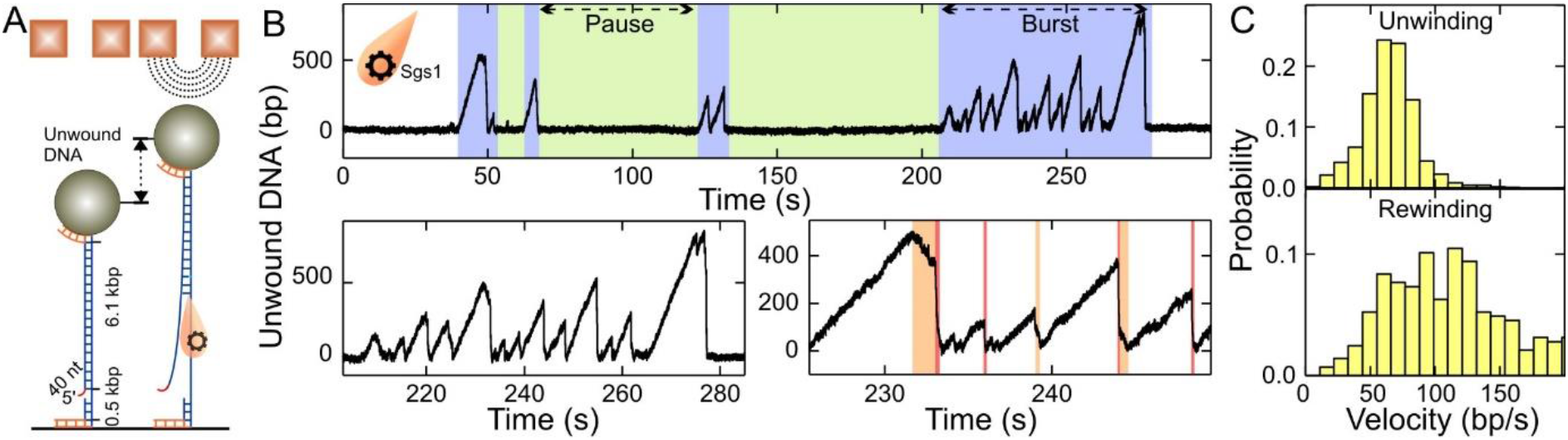
DNA duplex processing by Sgs1. A Sketch of the employed magnetic tweezers assay. B Observed dsDNA processing patterns of Sgs1, including an overview over consecutive bursts and pauses (upper panel) as well as detailed views into individual bursts containing multiple unwinding events (lower panels). A typical unwinding event of Sgs1 starts with slow, gradual unwinding of the dsDNA followed by DNA rehybridization, that can be almost instant (76% of events, see pink sections in lower right panel) or contain slow rewinding sections (24% of cases, see orange sections in lower right panel). C Histograms of the observed unwinding and rewinding velocities for Sgs1. The mean unwinding rate was 65±2 bp/s (*N* = 899). The mean rewinding rate was 115±6 bp/s (*N* = 287). Data information: In C, data are presented as mean ± SEM.

When adding Sgs1 in the presence of ATP into the fluidic cell, DNA unwinding was seen as a gradual increase of the DNA length due to a larger extension of single-stranded DNA compared to dsDNA at the applied forces (Fig 1B). Consistently, no DNA lengthening, i.e. no unwinding, was observed when omitting ATP or protein in the reaction. The unwinding rates followed a Gaussian-like distribution with a mean of 65 ± 2 bp/s (Fig 1C, upper panel), indicating within error a unique unwinding rate for Sgs1. DNA unwinding was typically terminated by an abrupt DNA length decrease (pink sections in Fig 1B) that reflects rezipping, i.e. renaturation of the DNA duplex. Occasionally, the DNA rezipping contained short sections of a slow DNA length decrease (24% of the DNA closing events, see orange sections in Fig 1B, lower panel). Since the rate of these sections was approximately constant and was on the order of magnitude of the unwinding rate, we attribute these sections to helicase-driven DNA rewinding. We believe that Sgs1 is in these cases translocating on the opposite ssDNA strand away from the Y-junction, and thus limiting the rehybridization to the ssDNA translocation rate of the helicase. The single unwinding-rezipping events typically occurred in bursts comprising several individual events followed by long pauses (Fig 1B, upper panel). This indicated that a burst was likely initiated by the binding of a single Sgs1 unit (a molecule or a complex), which subsequently originated all events of the burst until the protein finally dissociated. Such behavior is similar to that observed for BLM [29], WRN [30] and the likely BLM-homologue from *Arabidopsis thaliana*, AtRecQ2 [31]. For the latter it has been shown that the transition between unwinding and rezipping most likely involves strand switching that brings the enzyme in a more loosely bound state since it lacks the DNA junction in its wake [31]. The lowered affinity predominantly causes fast rezipping, in which the enzyme is pushed by the rehybridizing junction, but occasionally slowly rewinds the DNA due to ssDNA translocation. New DNA unwinding needs to be reinitiated by an additional strand switch. Due to the functional similarities between Sgs1, BLM and AtRecQ2 [21,32], we suggest that Sgs1 also undergoes cycles of strand switches during repetitive DNA unwinding-rezipping events, suggesting that this is a conserved characteristic of RecQ helicases.

### DNA unwinding by Sgs1 in presence of RPA

To systematically probe the influence of protein partners on the behavior of Sgs1, we first studied its DNA unwinding capacity in the presence of RPA. Compared to Sgs1 alone, the unwinding-rezipping events appeared significantly altered when adding only 20 nM of the single-stranded DNA binding protein (Fig 2A). No fast DNA rezipping events were observed anymore, such that gradual DNA unwinding was exclusively followed by gradual DNA rewinding. Similarly to Sgs1 alone, events occurred in a repetitive, burst-like manner. The bursts were similarly separated by pauses (Fig 2A, upper panel), indicating that a single unwinding complex was likely driving the reaction.

**Fig 2.**
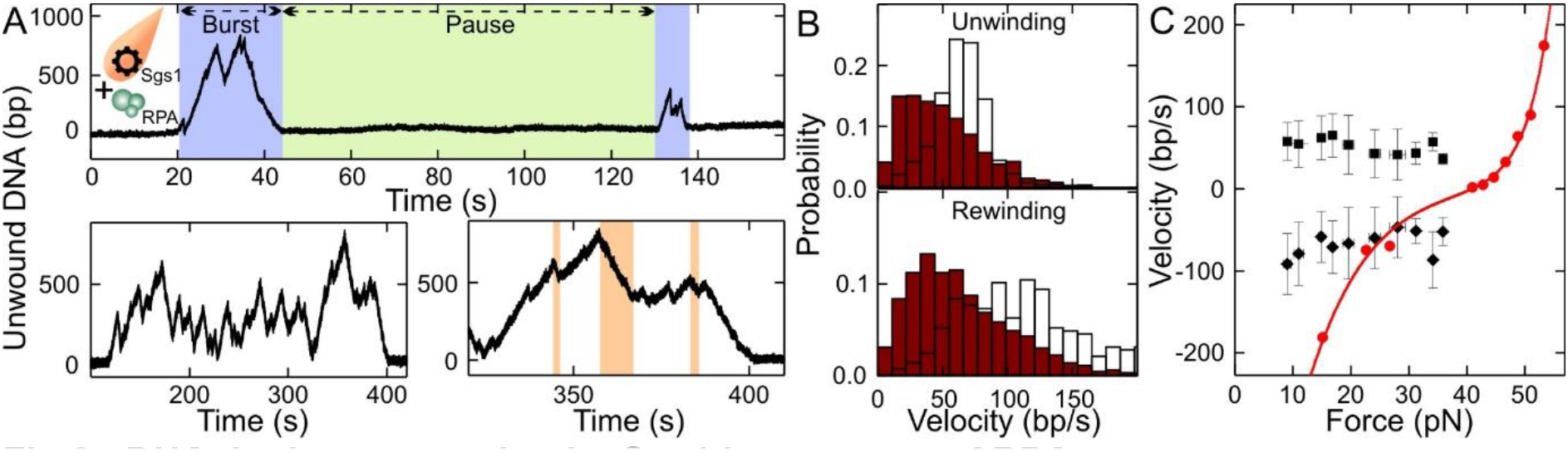
DNA duplex processing by Sgs1 in presence of RPA. A Observed dsDNA processing pattern of Sgs1 in presence of RPA including an overview over successive bursts and pauses (top panel) and detailed views of single bursts (bottom panels). Unwinding events are always followed by slow rewinding events (orange sections). B Histograms of the observed unwinding and rewinding velocities for Sgs1 in presence of RPA (brown bars). The mean unwinding rate was 51±3 bp/s (*N* = 2139). The mean rewinding rate was 66±3 bp/s (*N* = 2091). White bars show for comparison the distributions of unwinding and rewinding velocities for Sgs1 alone (taken from Fig 1C) for reference. We attribute the large rewinding rate in absence of RPA to an increased error in inferring the rewinding rate from the short rewinding sections. C Force-dependence of the Sgs1 unwinding (black squares) and rewinding rates (black diamonds) in presence of RPA (errors given as standard deviations). Red circles represent the force-dependent DNA opening and closure rates measured for RPA alone (taken from Ref. [33]. A fit to this data is shown as a solid red line.). Data information: In B, data are presented as mean ± SEM.

The observed slow rewinding could result from either a limited velocity of Sgs1 translocating along the RPA-coated ssDNA strand away from the junction or due to the limited rate at which RPA dissociated from the junction ends. To reveal whether active translocation or passive dissociation caused the slow rewinding, we carefully characterized the unwinding and rewinding velocities in a force range between 10 and 35 pN. No force dependence was detected for neither of the two processes (Fig 2C). The mean rates for unwinding and rewinding were 51 ± 3 bp/s and 66 ± 3 bp/s, i.e. rather similar (Fig 2B). This is in contrast to the expected rates at which RPA gets dissociated by a rezipping junction. Previous measurements found a strong exponential force dependence for such an RPA dissociation (see rate line in Fig 2C) [33]. Therefore, DNA unwinding as well as rewinding is an active process that is driven by the helicase rather than the association and dissociation of RPA.

While the mean unwinding velocity was only mildly reduced in the presence of RPA compared to its absence, the rate distribution markedly differed. In particular, the presence of RPA caused a strong skew of the distribution with a maximum at 30-40 bp/s (Fig 2B) in contrast to the Gaussian distributed rates in the absence of RPA. A similarly skewed distribution was observed for rewinding (Fig 2B). The shift of the rates towards lower values could be due to RPA acting as a roadblock in the way of Sgs1. However, this should affect unwinding to a much lower extent than rewinding, where Sgs1 has to displace RPA directly. We therefore attribute the skewed rate distributions to a direct interaction between Sgs1 and RPA, in which the RPA-bound form of Sgs1 has a slower and more variable unwinding/translocation rate. A similar skew was also observed at elevated RPA concentrations (50 nM) suggesting that Sgs1 populations that would differ in the number of bound RPA molecules are not the primary reason for the observed rate distributions. Overall our conclusions are consistent with previous reports, which demonstrated multiple RPA binding regions on Sgs1 [25] such that different binding states of RPA could modulate the Sgs1 behavior.

### DNA processing by the combined activity of Sgs1 and Dna2

Next, we set out to investigate the full DNA end resection reaction in our setup that includes Sgs1, RPA and Dna2. First, we tested DNA unwinding by Dna2 in the presence of RPA but in the absence of Sgs1. The Dna2 unwinding activity requires a 5’ ssDNA flap as present in our DNA substrate (Fig 1A). For wt Dna2, the nucleolytic degradation of such a flap wins over initiation of unwinding, which effectively inhibits DNA unwinding by the helicase activity of the wt enzyme [13]. However, processive DNA unwinding was seen with the nuclease-dead Dna2 E675A mutant (Fig 3A, inset). In agreement with earlier investigations [13,14] as well as its barrel-shaped, DNA encircling structure [17], the DNA unwinding by Dna2 was completely unidirectional, i.e. no DNA rewinding was observed at all. The unwinding rates by Dna2 were highly variable between the individual enzyme molecules ranging from 15 to 160 bp/s [13].

**Fig 3.**
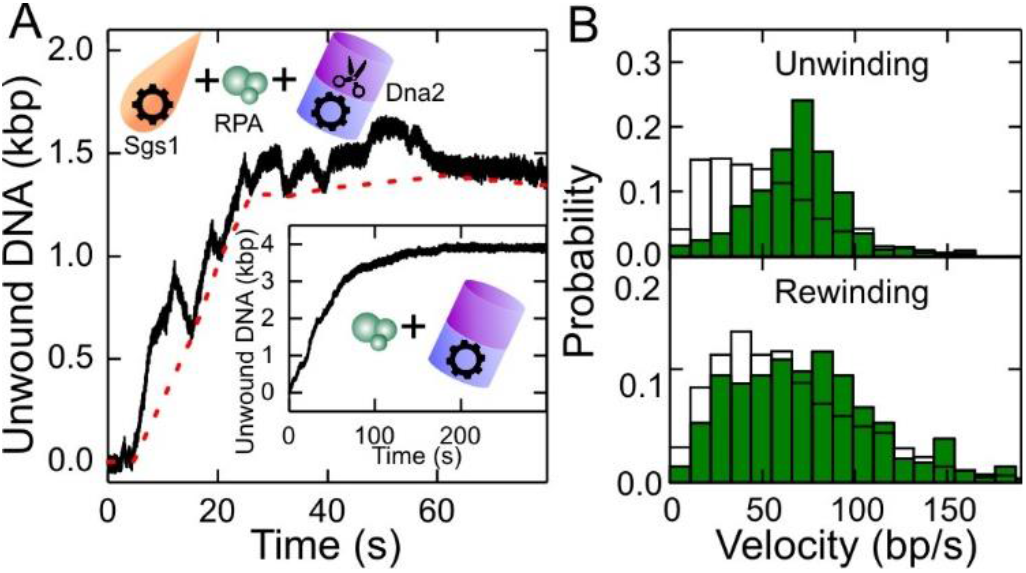
DNA duplex processing by the combined activity of Sgs1, Dna2 and RPA. A Typical dsDNA processing event. The dashed red line connects the minima at the end of the rewinding sections (see text for details). The inset shows a DNA unwinding event in presence of nuclease-dead Dna2 E675A and RPA. B Histograms of the observed unwinding and rewinding rates (green bars) with mean values of 69±3 bp/s (*N* = 365) and 74±5 bp/s (*N* = 282), respectively. White bars show for comparison the unwinding and rewinding velocities of Sgs1 in the presence of RPA (taken from Fig 2B). Data information: In B, data are presented as mean ± SEM.

After confirming the activity of Dna2 alone, we studied the minimal reconstituted DNA end resection reaction in the presence of Sgs1, wt Dna2 and RPA monitored by DNA unwinding. Notably, in contrast to wt Dna2 in the absence of Sgs1, significant DNA unwinding over kpb distances was observed (Fig 3A). The observed trajectories exhibited a combination of the unwinding patterns of both helicases investigated individually. Particularly, the typical Sgs1 patterns of alternating sections of DNA unwinding and rewinding were observed. However, we observed a gradual increase of baseline DNA unwinding, i.e. Sgs1 was not capable to fully rewind the DNA anymore (Fig 3A). This indicated that Dna2 was progressively moving along the 5’ DNA end, which was limiting the translocation of Sgs1 along this strand when going backwards. This may be due to Dna2 physically blocking DNA reannealing, or DNA degradation. In this case, the approximate translocation/unwinding/DNA degradation distance was obtained by connecting the lower turning points of Sgs1 in the trajectories (see dashed lines in Fig 3A and Fig EV1A). In addition to wt Dna2, we examined the nuclease-dead (Dna2 E675A) as well as the helicase-dead (Dna2 K1080E) Dna2 mutants in combination with Sgs1 and RPA (Fig EV1B, EV1C). For both Dna2 mutants we observed a similar progressive DNA unwinding overlayed by short Sgs1 unwinding-rewinding cycles as for wt Dna2. The data for the nuclease-dead variant suggested that already the helicase motor alone can act as a sufficiently strong road block to stop DNA rewinding by Sgs1. Interestingly, upon removing the excess enzyme from the flow cell and additionally challenging the complex with 3 M NaCl after an unwinding reaction, the DNA remained unwound also for the nuclease dead mutant (i.e. no rezipping occurred). This corroborates that Dna2 forms an irreversible road block on the 5’ end as it encircles the DNA strand and thus cannot dissociate. The observation that helicase-dead Dna2 also functions as a road block for DNA rewinding by Sgs1 indicates the progressive degradation of the 5’ DNA end by its nuclease domain. This confirms that the Dna2 helicase activity is not required for end resection [*15*]. In this case the nuclease domain is thought to employ an electrostatic ratchet mechanism [*17*,*34*] to progressively degrade the 5’ DNA end.

When inspecting the rates of the Sgs1 unwinding and rewinding cycles of the end resection reaction containing wt Dna2, we found a Gaussian-like distribution for both unwinding and rewinding with significantly reduced skews compared to Sgs1 in the presence of RPA (Fig 3B, Fig EV2). This suggests that Sgs1 is likely directly interacting with Dna2 in the end resection reaction both during unwinding and rewinding such that it can alleviate the inhibitory activity of RPA on the Sgs1 unwinding velocity. Since rewinding included only slow events, we conclude that RPA is still interacting with Sgs1, such that a ternary complex of the three different proteins is formed. Overall these data show that both proteins act simultaneously during the end resection reaction, and unwind/translocate DNA at different velocities. The modulation of the velocity of Sgs1 by Dna2 suggests that there is some degree of coupling/coordination in addition to complex formation between the two enzymes.

### DNA unwinding by Sgs1 in complex with Top3-Rmi1

It is well established that Top3-Rmi1 forms a complex with Sgs1 via interaction at the far end of the N-terminus of Sgs1 [35]. Top3-Rmi1 cooperates with Sgs1 in DSB end resection [36,37] as well as in the dissolution of double Holliday junctions [37,38]. Generally, Top3-Rmi1 is known to stimulate the rates of Sgs1 unwinding and DNA resection in biochemical assays [19], however the underlying mechanism remains unclear. In order to study how the complex formation with Top3-Rmi1 affects the activity of Sgs1, we conducted a set of experiments using our magnetic tweezers assay.

We first tested the activity of Top3-Rmi1 alone on our flapped DNA substrate. Surprisingly, we observed step-wise increases of the DNA length of ~8 nm length (Fig EV3B). This activity required the presence of a single stranded region on the DNA substrate. We attributed it to the ssDNA cleavage activity of the type 1A topoisomerase Top3 [39]. Upon DNA cleavage Top3 adopts an open form, covalently attached to 5’-end of cut strand [40,41]. This can result in the formation of an elongated Top3-DNA chain (Fig EV3A).

When testing Sgs1 and Top3-Rmi1 together, we observed the step-wise length increases due to the likely formation of Top3 bridges, as well as the typical saw tooth-like pattern from DNA unwinding by Sgs1 (Fig EV3D). Sgs1 unwinding could be clearly discriminated from Top3-Rmi1 bridge formation and occurred in a burst-like manner. Individual unwinding events comprised a gradual unwinding and typically an abrupt rezipping of all unwound DNA as seen for Sgs1 alone (Fig 4A). The complex formation with Top3-Rmi1 reduced the fraction of partial rewinding events from 24% to 5%. The unwinding velocities were Gaussian-like distributed and were more than 30% faster for the Sgs1-Top3 complex as compared to Sgs1 alone (86 ± 3 bp/s instead of 65 ± 2 bp/s, see Fig 4B).

**Fig 4.**
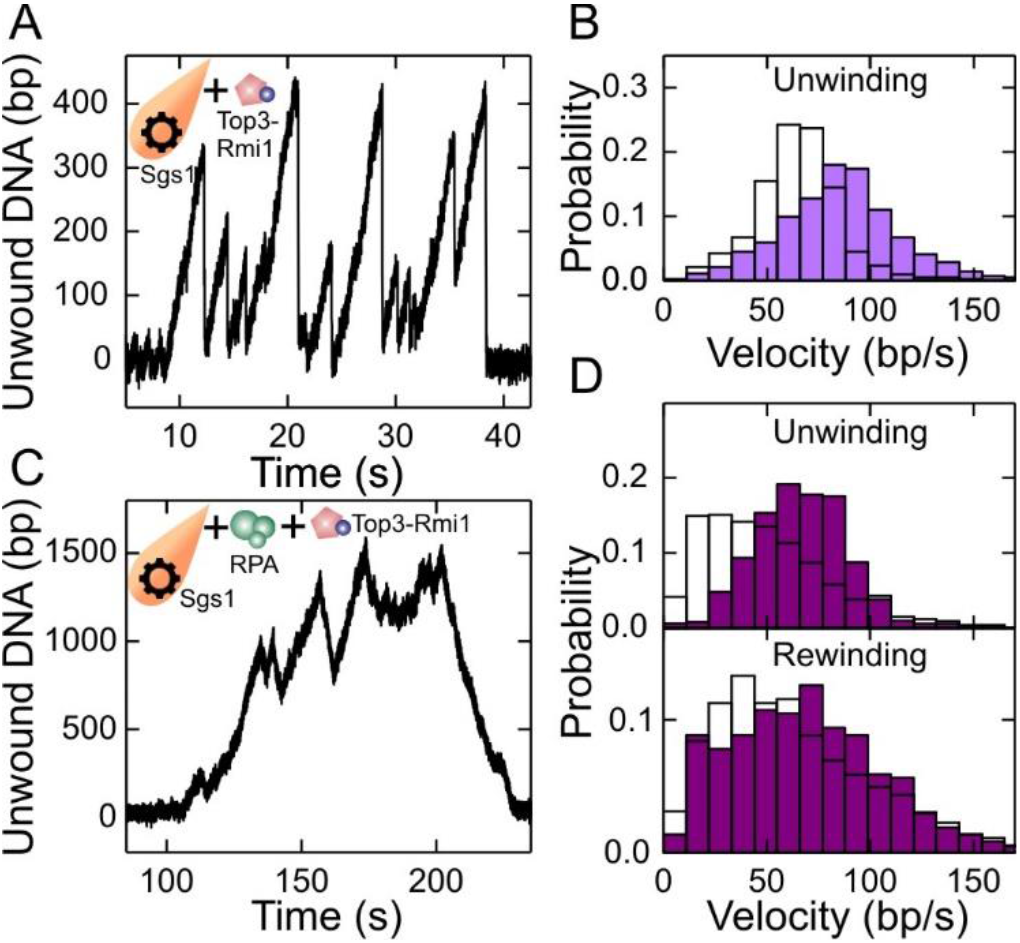
DNA duplex processing by Sgs1 in presence of Top3-Rmi1. A dsDNA processing events observed for Sgs1 in presence of Top3-Rmi1. Typical events consist of periods with slow gradual unwinding typically followed by instant DNA rezipping as seen also for Sgs1 alone. B Histogram of the unwinding rate of Sgs1 in presence of Top3-Rmi1 (violet bars) with a mean rate of 86 ± 3 bp/s (*N* = 1101). For comparison, the distribution of unwinding rates for Sgs1 alone is depicted with white bars (taken from Fig 1C). C dsDNA processing by Sgs1 in presence of Top3-Rmi1 and RPA. In contrast to the absence of RPA, DNA closure is seen as a slow rewinding. D Histograms of unwinding and rewinding rates of Sgs1 in presence of Top3-Rmi1 and RPA (purple bars). The mean rates for unwinding and rewinding are 66 ± 2 bp/s (*N* = 502) and 70 ± 4 bp/s (*N* = 369), respectively. For comparison, the velocity distributions for Sgs1 in presence of RPA are shown as white bars (taken from Fig 2B). Data information: In B and D data are presented as mean ± SEM.

When studying DNA unwinding by the Sgs1-Top3-Rmi1 complex in the presence of RPA no sudden length increases due to the formation of Top3 bridges were observed. This indicates that RPA protects ssDNA from the cleavage activity of Top3-Rmi1. Gradual DNA unwinding was always followed by a gradual DNA rewinding as also seen for Sgs1 and RPA alone (Fig 4C). Unwinding and rewinding velocities were higher than for Sgs1 and RPA alone by 30% and 5%, respectively, which suggests that Top3-Rmi1 generally accelerates the motion of Sgs1 on DNA. Most importantly, the distribution of the unwinding and rewinding velocities was Gaussian-like and significantly less skewed compared to Sgs1 and RPA alone (Fig 4D, Fig EV2). This provides an independent control for complex formation between Sgs1 and Top3-Rmi1, which is similar to the reactions with Sgs1 and Dna2, for which an unskewed velocity distribution was also obtained.

Top3-Rmi1 is known to stimulate the activity of Sgs1 in particular at elevated salt concentrations [19]. To test this in our experiments, we challenged DNA unwinding by Sgs1 with “high salt” conditions (100 mM NaCl, 5 mM Mg^2+^) being close to physiological ionic strength. Independent of the presence of Top3-Rmi1, no DNA unwinding by Sgs1 was observed in the absence of RPA. When supplementing Sgs1 with RPA similar unwinding-rewinding events as found for our standard reaction condition were observed (Fig EV4A). At 20 nM RPA, the velocity distribution of Sgs1 was however little skewed (Fig EV4B, gray bars). Complexes between RecQ helicases and RPA [42,43] are known to be rather stable. However, electrostatic interactions can become screened at elevated salt concentrations [44], which can effectively increase the dissociation constant of the interaction. In agreement with this hypothesis, the Sgs1 unwinding velocity was found to be highly skewed in the presence of 200 nM RPA (Fig EV4B, dark blue bars). It also did not exhibit any significant force dependence (Fig EV4C). Addition of Top3-Rmi1 to the reaction (Fig EV4D) reduced the skew in the velocity distribution, indicating complex formation with Sgs1. Furthermore, the mean velocity in the presence of Top3-Rmi1 was increased by 23%. Given that RPA is an abundant protein, these results indicate that the observed velocity modulations by addition of RPA and Top3-Rmi1 are relevant at physiological salt concentrations and that Top3-Rmi1 serves as a general accelerator of the Sgs1 motor activity.

### RPA as important determinant of Sgs1 unwinding activity

So far we analyzed the unwinding and rewinding velocities by Sgs1. However, in addition to an overall DNA unwinding, observed in vivo or in a test tube, other parameters of the whole reaction - including the rate of recruitment to the DNA template, the burst duration and the processivity of unidirectional unwinding and rewinding - play an important role.

When analyzing the pause durations between individual bursts (see Fig 1B, 2B top panels), which are the inverse of the recruitment rate (Fig 5A, Fig EV5A), we found longer pauses at high salt compared to standard reaction conditions. Overall, we did not see significant differences between the absence or the presence of the cofactors RPA and Top3-Rmi1. Thus, at standard reaction conditions neither RPA nor Top3-Rmi1 contribute to the recruitment of Sgs1. At high salt conditions RPA was essential for activity, i.e. recruitment, while Top3-Rmi1 had only little influence. The influence of cofactors was different when analyzing the mean duration of the full unwinding bursts comprising many individual unwinding events (Fig 1B, 2B top panel). In the absence of RPA burst durations were only 32 s on average but could exceed 200 s in presence of RPA (Fig 5A, Fig EV5A). Top3-Rmi1 had little influence on the burst duration. Thus, RPA is a major determinant for the affinity of Sgs1 to the unwinding fork in agreement with the formation of a Sgs1-RPA complex.

**Fig 5.**
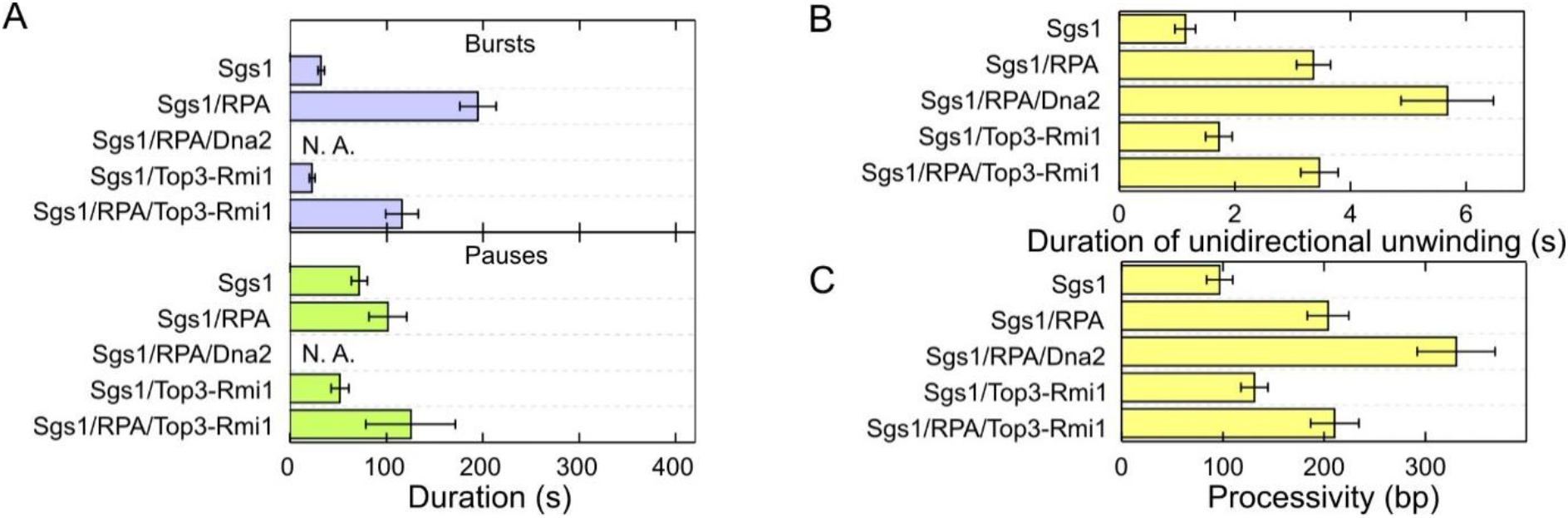
Directionality, processivity and initiation of DNA processing events for the different protein combinations. A Mean duration of bursts (violet bars) and pauses (green bars) for the different enzyme combinations. B Mean duration of a continuous unwinding event. C Mean processivity of unidirectional DNA unwinding.

Finally, we analyzed the processivity of DNA unwinding by Sgs1. This can be either presented as the time during which Sgs1 is unidirectional unwinding without direction reversal (Fig 5B) or as a number of base pairs that are unwound during this time interval (Fig 5C). The addition of RPA to the reaction had the most significant influence, which increased the duration of continuous unwinding 3 to 6-fold and the processivity 2 to 3.5-fold. Top3-Rmi1 increased the processivity more moderately and only at normal reaction conditions. Altogether, these data reveal that RPA modulates the recruitment at high salt, the affinity to the unwinding fork as well as the processivity of DNA unwinding by Sgs1.

## Discussion

In this study we characterized how the unwinding of DNA by the yeast RecQ family helicase Sgs1 is modulated by protein cofactors that cooperate with it during DNA end resection. We showed that a single Sgs1 protein complex can drive processive DNA unwinding over hundreds of base pairs (Fig 1). DNA unwinding by Sgs1 was found to be highly dynamic, involving many repetitive unwinding events separated by either rapid DNA rezipping or slower DNA rewinding. Sgs1 shares this highly dynamic activity pattern with other RecQ family helicases from prokaryotes [45,46] and eukaryotes [29–31]. It is thought that the switching between unwinding and rewinding involves repeated strand switching events to allow direction reversals of the helicase [31]. In the absence of any cofactor the unwinding velocity of Sgs1 had a narrow distribution. In the presence of RPA the distribution became rather broad and strongly skewed. Furthermore, only slow DNA rewinding due to active translocation by Sgs1 rather than fast rezipping was observed. We attributed this changed behavior to the formation of a Sgs1-RPA complex. The broad distribution of the Sgs1 velocities with RPA may be due to multiple binding states of RPA that modulate the Sgs1 behavior in a different manner [25].

When reconstituting the DNA end resection reaction by combining Sgs1, RPA and Dna2, we observed that Sgs1 continued its dynamic DNA unwinding-rewinding activity including frequent direction reversals. In addition, the presence of Dna2 promoted a progressive overall unwinding, i.e. rewinding events did not succeed to close the full DNA duplex, but rather terminated away from the original flap position at a distance that increased with time (Fig 3, Fig EV1). This observed behavior is in agreement with the stringent unidirectional DNA unwinding of Dna2 [13,14] (see inset in Fig 3), combined with progressive degradation of the DNA 5’ end. Thus, the unwinding activity of Sgs1 and the unwinding/degradation activity of Dna2 occur at different velocities. Interestingly, when analyzing the Sgs1 velocities during DNA end resection, we observed that the distributions lost the pronounced skew observed in presence of RPA. This indicates that the unwinding activity of Sgs1 is to some degree coupled to Dna2 and that Sgs1 directly interacts with Dna2 by forming a complex. This is in agreement with previous biochemical data that revealed that Sgs1 and Dna2 can directly physically interact with each other [19]. The direct interaction appears to alleviate the inhibitory effect of RPA on the unwinding velocity, either by disrupting Sgs1-RPA contacts or by allosteric means. Since Dna2 requires RPA for correct loading onto the 5’ end [17] and since the rewinding events of Sgs1 were exclusively slow during end resection, we propose that RPA is still part of the formed DNA end resection machinery, forming a ternary complex.

Since only Sgs1 can unwind dsDNA, and the average speed of the Sgs1 motor is about two times higher compared to that of Dna2 [13], Sgs1 is the main factor for DNA unwinding, while Dna2 is trailing behind on the 5’ ssDNA end. Dna2 movement along the unwound ssDNA is powered by its motor activity [15,16]. Thus, a loop has to form in front of Dna2 (Fig 6). A similar loop formation has been found for the prokaryotic RecBCD complex [47], indicating that similar mechanisms can also exist in eukaryotic cells. A loop forming ahead of Dna2 would allow the binding of RPA to the unwound DNA behind Sgs1, which was shown to specifically promote degradation of the 5’-terminated strand by Dna2 [19,28]. A loop forming ahead of Dna2 thus explains how the regulatory function of RPA can be achieved. Next, Sgs1 occasionally switches strands and actively rewinds DNA, thus backtracking towards the slower moving Dna2 molecule. When Sgs1 encounters Dna2 it switches back again to DNA unwinding. What can be the reason for such a switching behavior? We hypothesize that the strand switching activity serves to limit the DNA unwinding by Sgs1, such that the ssDNA loop is not extensively long and prone to unscheduled cleavage. Furthermore, Dna2 is sensitive to obstacles on DNA such as secondary structures or protein blocks, which stall DNA degradation [13,48]. Sgs1, moving periodically toward Dna2, might help resolve these structures. Finally, Sgs1 has a function to promote the dissolution of double Holliday junctions in the late stage of the canonical DSB repair pathway [49], which is separate from its role in DNA end resection. To do so, its needs to migrate, in conjunction with Top3-Rmi1, the two Holliday junctions toward each other. However, it was not apparent how the direction of the junction migration by Sgs1 is determined, as only convergent migration (i.e. the migration of the junctions toward each other) dissolves the entangled chromosomes [49]. It has been speculated that chromatin may serve as a barrier to block junction migration in the “wrong” direction. This, coupled with the random switching of Sgs1 movement, would result in a “random walk” mechanism of junction migration, which would ultimately lead to convergence of the both junctions. Thus, the switching behavior of Sgs1 may be also relevant for processes separate from DNA end resection.

**Fig 6.**
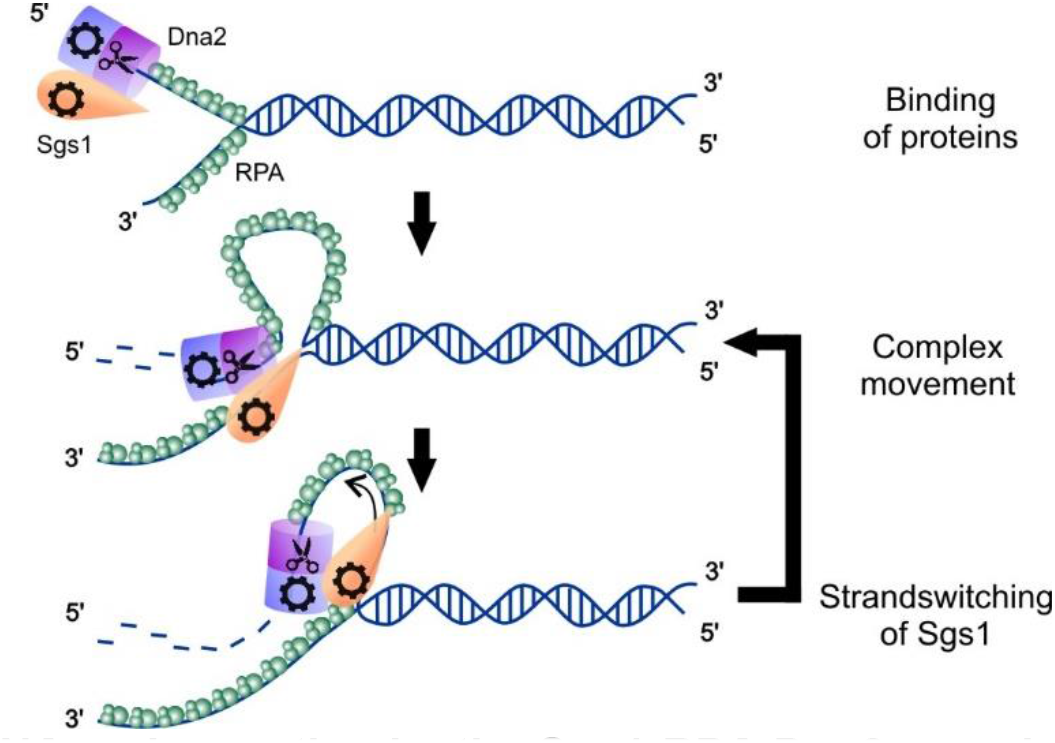
Model for DNA end resection by the Sgs1-RPA-Dna2 complex. A Sgs1-Dna2 complex binds first to a 5’ DNA overhang assisted by RPA. At the DNA junction both helicases start to move unidirectionally on either of the strands. Due to the faster movement of Sgs1, this helicase powers the unwinding of the DNA duplex and causes the formation of a ssDNA loop forms in front of Dna2. Upon an occasional strand switch of Sgs1, the protein rewinds the DNA and moves towards Dna2. This leads to shortening of the loop. Upon encounter of Dna2, Sgs1 switches back to the original strand to power further unwinding.

We also determined the effect of the known interaction partner Top3-Rmi1 on DNA unwinding by Sgs1. Most importantly, we found that in the presence of RPA, the histograms of the Sgs1 unwinding and rewinding velocities exhibited significantly reduced skews (Fig 4D) compared to the presence of RPA only. Similarly, the interaction with Top3-Rmi1 appeared to alleviate the inhibitory effect of RPA on the unwinding velocity. Additionally, we observed that the velocity of Sgs1 was increased in the presence of Top3-Rmi1 regardless of RPA (Fig 4B, D) or the ionic strength (Fig EV4E). An increased unwinding/rewinding velocity thus explains the mechanism underlying the previously observed stimulation of Sgs1 by Top3-Rmi1 [19]. Since in the presence of RPA DNA rewinding of the Sgs1-Top3-Rmi1 complex was always slow, we conclude that Sgs1 is simultaneously interacting with RPA and Top3-Rmi1. Notably, we observed that under the applied force, Top3-Rmi1 can be trapped on ssDNA in the so-called open gate configuration (Fig EV3A), which seems to be a general property of type IA topoisomerases [41]. This conformational trapping may however be less relevant in vivo since it is abolished by the presence of RPA, which limits the access of Top3-Rmi1 to ssDNA. The physiological relevance of this ssDNA cleavage activity of Top3-Rmi1 may thus be limited.

As Top3-Rmi1, also RPA appears to be an important regulator of Sgs1 activity. It is essential for the recruitment of Sgs1 at high ionic strength, which is similar to *E. coli* RecQ whose initiation is supported by SSB [50]. Furthermore, RPA slows the rewinding of DNA by Sgs1 (Fig 2A) and increases the processivity of Sgs1 for unidirectional unwinding (Fig 5C). Finally, RPA increases the duration of unwinding bursts, i.e. it stabilizes the interaction of Sgs1 with the DNA substrate (Fig 5B). Altogether, these results indicate that RPA ensures that DNA is kept in a partially unwound state for longer periods of time, which may promote the end resection process.

Altogether, our data show that the various Sgs1 protein partners lead to surprisingly diverse modulations of the Sgs1 activity. We believe that these modulations allow fine tuning of DNA unwinding by Sgs1, which helps it to tackle its diverse functions to promote genome stability.

## Methods

### Recombinant proteins

Sgs1, RPA, Dna2 from *Saccharomyces cerevisiae* and their mutants were expressed and purified as described previously [13,19,51]. In short, MBP-Sgs1 was expressed using pFB-MBP-Sgs1-His vector and the Bac-to-Bac baculovirus expression system in *Sf*9 cells. The protein was first bound to amylose resin (New England Biolabs). Afterwards, the protein was treated with PreScission protease to remove the MBP-tag. Next, Sgs1 was bound to Bio-Rex70 resin and Ni^2+-^NTA-agarose, eluted and dialysed [11]. Yeast RPA was expressed in *Escherichia coli* from p11d-scRPA vector (a kind gift from M. Wold, University of Iowa) and purified as described for the human recombinant RPA [52]. Wild type Dna2 and its mutants were expressed from altered pGAL:DNA2 vector containing N-terminal Flag and HA tags as well as C-terminal His6 tag, in *S. cerevisiae* strain WDH668 [53]. Dna2 was purified by affinity chromatography using Ni^2+^-NTA-agarose (Qiagen) and M2 anti-FLAG affinity resin (Sigma), washed and eluted with buffer containing 3xFLAG Peptide (Sigma) [13].

### DNA substrate

The DNA construct for the magnetic tweezers experiments containing the 40 nt flap (see Fig 1A) was prepared as previously described [13,54]. The main DNA fragment of 6.6 kbp in length was excised from plasmid pNLrep [55] using the restriction enzymes BamHI and BsrGI. It was simultaneously digested with the nicking enzyme Nt.BbvCI to produce a 63 nt gap at an engineered site containing 5 consecutive, 15 nt spaced Nt.BbvCI sites. The gap was located approximately 0.5 kbp from the BamHI–cut end of the fragment. 63 nt of the gap were filled by hybridizing a 25 nt DNA oligomer that carried an additional 40 nt polythimidine tail on its 5’-end that served as the flap. In a subsequent ligation reaction the oligomer was ligated at its 3’ end inside the gap. Furthermore, 600 bp DNA handles carrying either multiple biotin or digoxigenin modifications were attached at either end. The handle duplexes were produced by PCR in the presence biotin and digoxigenin modified nucleotides and digested with BsrGI and BamHI, respectively.

### Magnetic tweezers experiments

Single-molecule experiments were carried out in a custom-made magnetic tweezers setup [56,57] at room temperature. Fluidic cells were prepared from two coverslips (Menzel, Braunschweig, Germany) and a Parafilm (Bemis, Oshkosh, USA) spacer. The bottom coverslip was previously coated by spin-coating using a 1% solution of polystyrene in toluene (Sigma-Aldrich, St. Louis, USA). To allow specific DNA tethering in the fluidic cell, anti-digoxigenin antibodies (Roche, Penyberg, Germany) were adsorbed to the polystyrene layer overnight from a 50 mg/ml anti-digoxigenin in standard aqueous phosphate buffered saline (PBS) solution. Subsequently, the fluidic cell was incubated with 10 mg/ml bovine serum albumin (BSA, New England Biolabs, Ipswich, USA) to prevent non-specific surface binding. 3 μm latex beads (Life Technologies, Darmstadt, Germany) serving as reference particles and 2.8 μm streptavidin-coated magnetic beads (Life Technologies, Darmstadt, Germany) with prebound DNA molecules were flushed into the flowcell. The beads and the DNA was allowed to bind to the surface and subsequently unbound particles were removed by washing the chamber with phosphate buffered saline (PBS). Lowering the magnets, allowed to stretch and to identify bead-tethered DNA molecules. The measurements were then performed at 300 Hz using videomicroscopy and real-time GPU accelerated image analysis [56]. One measurement usually was performed with 15-25 molecules at a time. Magnetic forces were calibrated using fluctuation analysis [58]. Unless stated otherwise, the measurements were performed in reaction buffer (25 mM Tris-acetate pH 7.5, 2 mM magnesium acetate, 1 mM ATP, 1 mM DTT, 0.1 mg/ml BSA) using protein concentrations of 0.2 nM Sgs1, 20 nM yRPA, 0.4 nM Top3-Rmi1 and 5 nM of Dna2 or its variants.

### Data analysis

Analysis of the results was performed using custom-written MATLAB program [59]. Particularly, the unwinding and rewinding velocities were determined from fitting linear segments to periods of constant velocities of the recorded trajectories. For converting measured unwinding velocities in μm/s into unwinding rates in bp/s, a conversion factor was obtained from recording force-extension curves of bare DNA construct and RPA coated construct. Errors of obtained rates and times are given as standard error of the mean (S.E.M.) throughout.

## Acknowledgements

We greatly acknowledge stimulating discussions and experimental support by Felix Kemmerich, Marius Rutkauskas, Dominik Kauert and Pierre Aldag. This work was supported by a Consolidator grant of the European Research Council (724863) to R.S. and by the Swiss National Science Foundation (31003A_175444) and European Research Council grants (681630) to P.C.

## Author contributions

P.C. and R.S. designed the study. K.K., P.C. and R.S. wrote the manuscript. K.K., F.F. and P.D. carried out the measurements and analyzed the data. M.L. purified and characterized the employed proteins.

## Conflict of interest

The authors declare that they have no conflicts of interest.

