## Supplementary Information for "Competing interactions modulate the activity of Sgs1 during DNA end resection"

### Expanded View

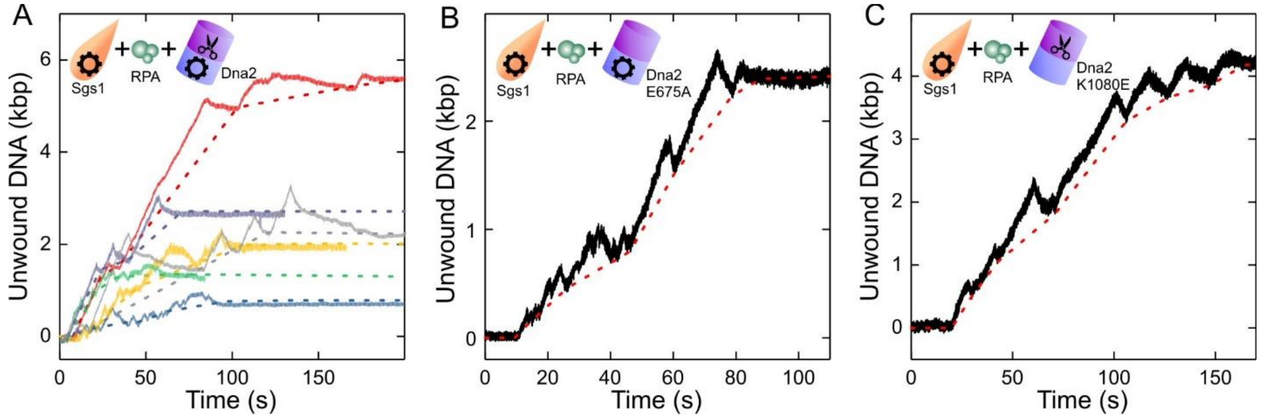


**Fig EV1 - DNA duplex processing by the DNA resection complex containing Sgs1, RPA and mutated Dna2.**

A Different dsDNA processing events in presence of Sgs1, Dna2 and RPA. Solid lines show the measured DNA unwinding, while dashed lines reveal the estimated Dna2 position.

B dsDNA processing event in presence of Sgs1, RPA and the nuclease-dead mutant Dna2 E675A. (c) dsDNA processing event in presence of Sgs1, RPA and the helicase-dead mutant Dna2 K1080E.

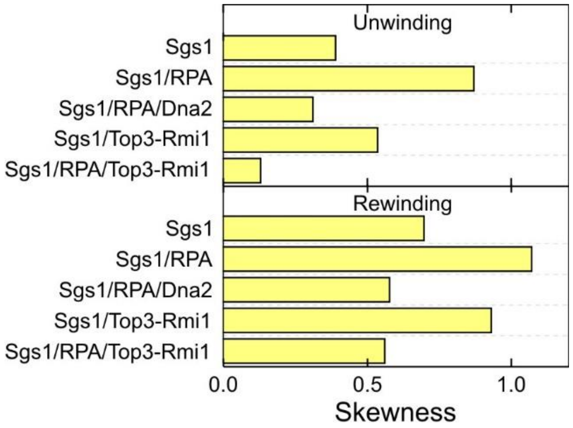


**Fig EV2 - Skewness of the velocity distributions for unwinding and rewinding for the different protein combinations.**
The skewness was calculated for the data shown in the velocity histograms in the main text.

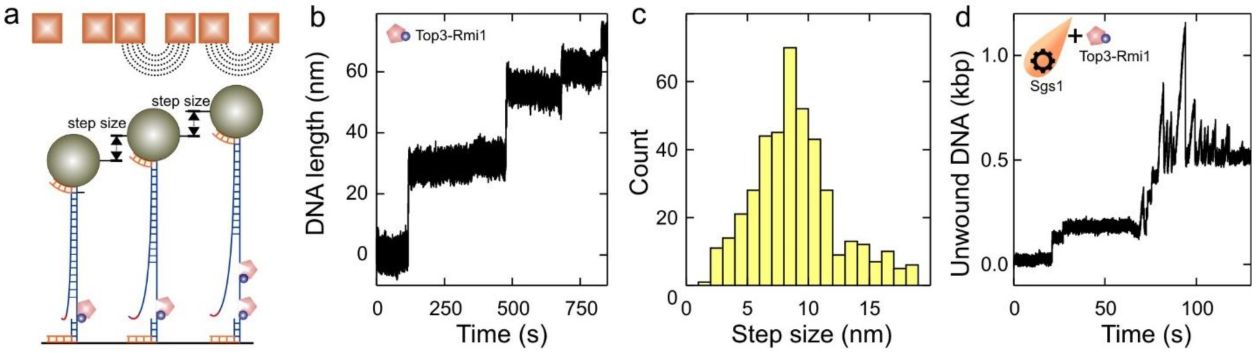


**Fig EV3 - Top3-Rmi1 cleavage activity of ssDNA.**

A Scheme illustrating the Top3-Rmi1 ssDNA cleavage activity observed in the magnetic tweezers assay.

B Stepwise DNA length increase seen in presence of Top3-Rmi1.

C Histogram of the size of the step-wise length increase having a mean of 8.6 ± 0.2 nm (*N*=433).

D dsDNA processing events observed for Sgs1 in presence of Top3-Rmi1. Step-wise DNA elongation due to ssDNA cleavage by Top3-Rmi is observed together with dsDNA unwinding-rezipping bursts by Sgs1.

Data information: In C, data are presented as mean ± SEM.

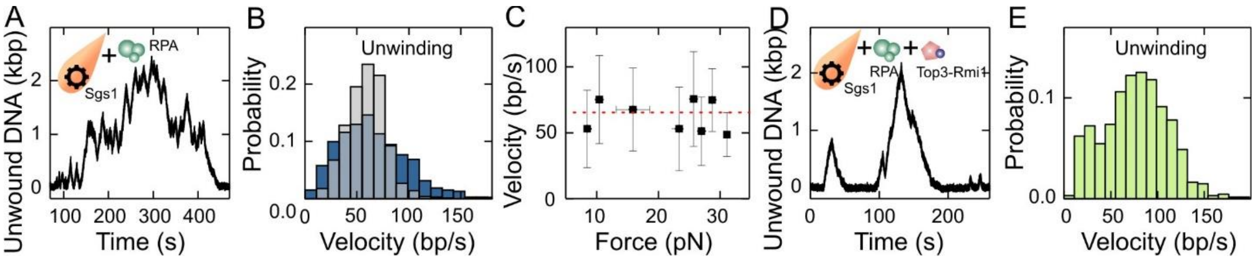


**Fig EV4 - DNA duplex processing by Sgs1 together with RPA and Top3-Rmi1 in high salt buffer.**

A dsDNA processing events observed for Sgs1 in presence of RPA in high salt buffer.

B Histograms of unwinding velocity of Sgs1 in presence of RPA in high salt buffer. Shown are the velocities in presence of 20 nM RPA (gray) and 200 nM RPA (dark blue). Mean unwinding velocities are v=59±2 bp/s (SEM, *N* = 860) and v=65±3 bp/s (*N*=1095) for 20 and 200 nM RPA, respectively.

C Force dependence of the Sgs1 unwinding (black squares) velocities in presence of 200 nM RPA. Red dashed lines represent mean values of the velocities, error bars represent the standard deviation of the mean.

D dsDNA processing events observed for Sgs1 in presence of RPA and Top3-Rmi1 in high salt buffer.

E Histogram of the unwinding velocity of Sgs1 in presence of RPA and Top3-Rmi1. The mean unwinding velocity was v=77±4 bp/s (*N*=678).

Data information: In B and D data are presented as mean ± SEM.

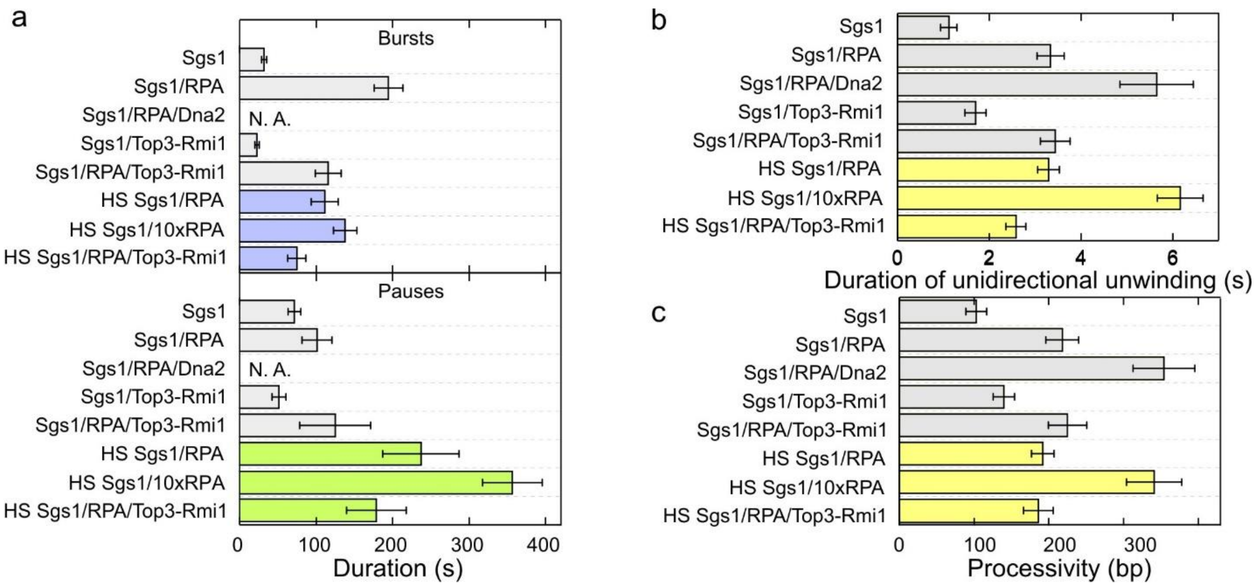


**Fig EV5 - Directionality, processivity and initiation of DNA processing events for the different protein combinations at high salt conditions.**

A Mean duration of bursts (violet bars) and pauses (green bars) for the different enzyme combinations.

B Mean duration of unidirectional unwinding. (d) Mean processivity of unidirectional unwinding by the enzyme(s).
